# Species-specific developmental timing is associated with global differences in protein stability in mouse and human

**DOI:** 10.1101/2019.12.29.889543

**Authors:** Teresa Rayon, Despina Stamataki, Ruben Perez-Carrasco, Lorena Garcia-Perez, Christopher Barrington, Manuela Melchionda, Katherine Exelby, Victor Tybulewicz, Elizabeth M. C. Fisher, James Briscoe

## Abstract

What determines the pace of embryonic development? Although many molecular mechanisms controlling developmental processes are evolutionarily conserved, the speed at which these operate can vary substantially between species. For example, the same genetic programme, comprising sequential changes in transcriptional states, governs the differentiation of motor neurons in mouse and human, but the tempo at which it operates differs between species. Using in vitro directed differentiation of embryonic stem cells to motor neurons, we show that the programme runs twice as fast in mouse as in human. We provide evidence that this is neither due to differences in signalling, nor the genomic sequence of genes or their regulatory elements. Instead, we find an approximately two-fold increase in protein stability and cell cycle duration in human cells compared to mouse. This can account for the slower pace of human development, indicating that global differences in key kinetic parameters play a major role in interspecies differences in developmental tempo.

## INTRODUCTION

The events of embryonic development take place in a stereotypic sequence and at a characteristic tempo (*1, 2*). Although the order and underlying molecular mechanisms are often indistinguishable between different species, the pace at which they progress can differ substantially. For example, compared to their rodent counterparts, neural progenitors in the primate cortex progress more slowly through a temporal sequence of neuronal subtype production (*3*). Moreover, in different species of primates, the duration of progenitor expansion differs, which at least partly accounts for differences in brain size (*4, 5*). Even in more evolutionary conserved regions of the central nervous system (CNS) there are differences in tempo. The specification of neuronal subtype identity in the vertebrate spinal cord involves a conserved and well-defined gene regulatory programme comprising a series of changes in transcriptional state as cells acquire specific identities and differentiate from neural progenitors to post-mitotic neurons (*6*). The pace of this process differs between species, despite the similarity in the regulatory programme and the structural and functional correspondence of the resulting spinal cords. For example, the differentiation of motor neurons (MNs), a prominent neuronal subtype of the spinal cord, takes less than a day in zebrafish, 3-4 days in mouse, but in humans it takes ~2 weeks (*7, 8*). Moreover, differences in developmental tempo are not confined to the CNS. The oscillatory gene expression that regulates the sequential formation of vertebrate body segments – the segmentation clock – has a period that ranges from ~30mins in zebrafish, to 2-3h in mouse, and 5-6h in human (*9–11*). What causes this developmental allochrony - interspecies differences in developmental tempo - is unclear.

To address this question, we set out to compare the generation of mouse and human MNs in the developing spinal cord. Progenitors of the spinal cord arise from the caudal lateral epiblast in the posterior of the elongating embryo and initially express the transcription factors (TFs) Pax6 and Irx3 (*12*). Exposure to Sonic Hedgehog (Shh), emanating from the underlying notochord, results in ventrally located progenitors expressing Nkx6.1 and Olig2. This downregulates Pax6 and Irx3 (*13*). Olig2 and Nkx6.1 expressing progenitors are termed pMNs and these either differentiate into post-mitotic MNs, which express a set of TFs including Mnx1 and Isl1, or transition into p3 progenitors that express Nkx2.2 (*14*). This gene regulatory network (GRN), in which Olig2 plays a pivotal role, repressing Irx3 and Pax6 and promoting the differentiation of MNs, is conserved across vertebrates (*15*).

We took advantage of the in vitro differentiation of MNs from mouse and human embryonic stem cells (ESCs) to investigate the pace of differentiation. We find that MN differentiation in vitro recapitulates species-specific global timescales observed in the embryos, lasting ~3 days in mouse and more than a week in human. We show that increased levels of signalling are unable to speed up the rate of differentiation of human cells. Moreover, by assaying the expression of a human gene, with its regulatory landscape, in a mouse context, we rule out the possibility that species differences in genomic sequence plays a major role in temporal scaling. Finally, we provide evidence that differences in cell cycle, and protein kinetics can explain the differences in tempo indicating global differences in the kinetics of molecular processes affect developmental tempo.

## RESULTS AND DISCUSSION

The characteristic spatial-temporal changes in gene expression responsible for neural tube development are well described and the regulatory interactions between the genes involved have been defined (*16*). Despite the conservation of the gene regulatory network across vertebrates, only limited analysis has been performed on the relevant stages of human development (*17, 18*). We thus performed immunostainings on mouse and human embryonic spinal cords at brachial levels at equivalent stages (*19*) to more accurately correlate the major developmental events of neural differentiation processes in vivo between mouse and human (Fig. 1A). At their maximum extents, the OLIG2-expressing pMN domains comprise a large proportion of ventral progenitors, occupying approximately 30% of the dorsal-ventral length of the neural tube in both mouse and human embryos (Fig. 1B). Over the following two days of mouse development, from E9.5 to E11.5, many post-mitotic MNs differentiate (Fig. 1C) resulting in a marked reduction in the size of the pMN domain (Fig. 1B), despite the continued proliferation of the progenitors (*8*). By contrast, the pace of development is noticeably slower in human embryos. At Carnegie Stage (CS) 11 the pMN occupies a similar proportion of the human neural tube as the pMN in E9.0 mouse embryos. During the following 1-2 weeks of development (CS13-17, Fig. 1B), the size of the pMN decreases as MNs accumulate (Fig. 1C), but the rate of these change is slower than seen in mouse. MN production decreases at ~E11.5 in mouse and CS15-17 in human and glial progenitors, co-expressing SOX9 and NFIA, begin to arise in both species (Fig. 1D). Together, the data indicate an equivalent progression in neural tube development of mouse and human that lasts around 3 days in mouse and over a week in human (Fig. 1A).

**Figure 1.**
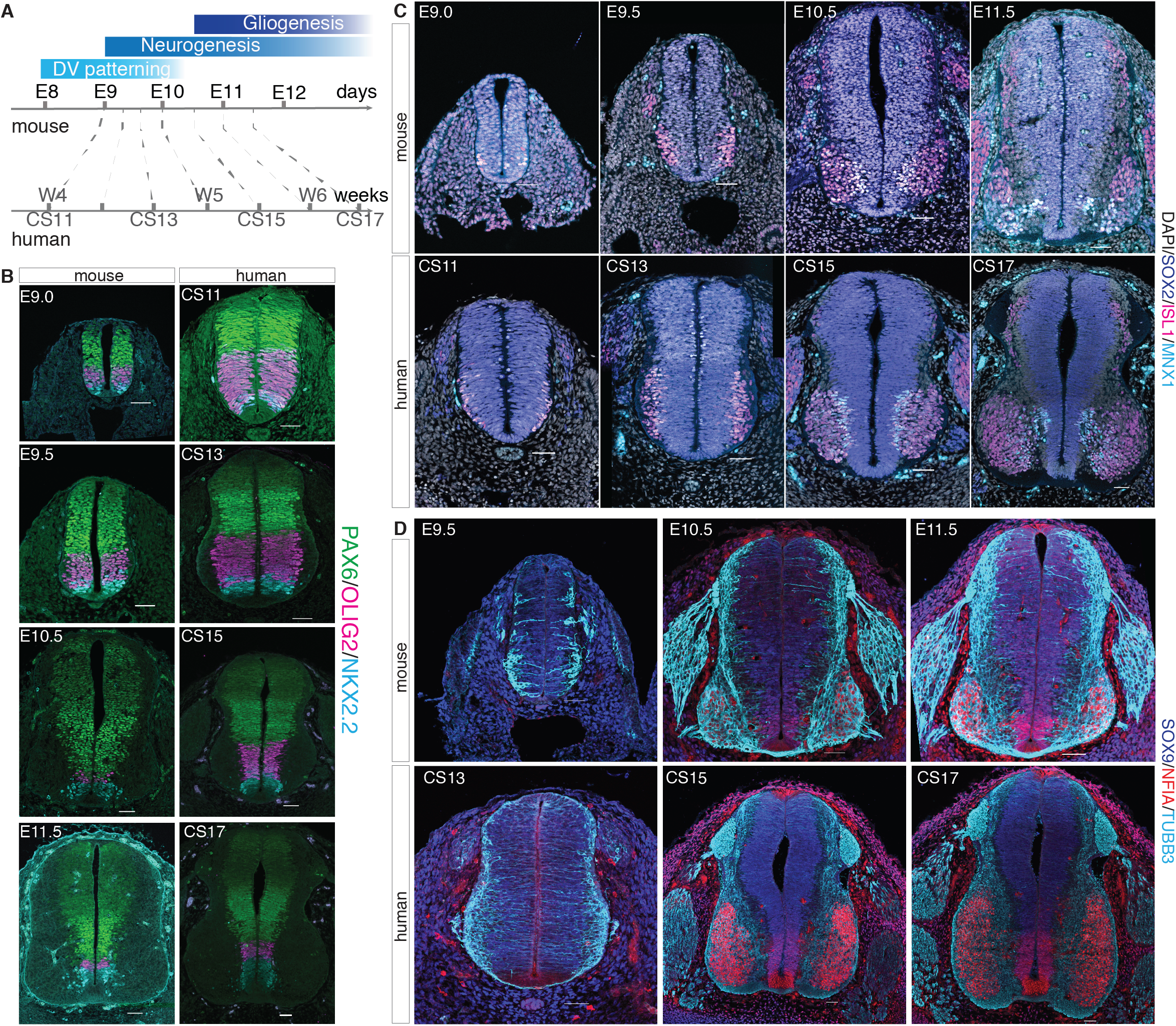
Comparison of neural tube development in mouse and human embryos. (**A**) Scheme matching mouse and human neural tube development according to neural patterning and differentiation. (**B-D**) Immunofluorescence in transverse sections of mouse and human neural tubes at shoulder levels from E9.0 to E11.5 in mouse and CS11 to CS17 in human embryos. (**B**) Expression of progenitor markers PAX6 (green), OLIG2 (magenta) and NKX2.2 (cyan). (**C**) Pan-neural progenitor marker Sox2 (blue), and motor-neuron markers ISL1 (magenta) and HB9/MNX1 (cyan) at neurogenic stages. (**D**) Early ventral expression of gliogenic markers NFIA (red) and SOX9 (blue) in the neural tube can be detected from E10.5 in mouse and CS15 in human. NFIA also labels neurons, as indicated by TUBB3 (cyan) staining. Scale bars = 50 μm.

We set out to examine whether interspecies tempo differences were preserved in vitro. Methods for the differentiation of MNs from ESCs, which mimic in vivo developmental mechanisms, have been established for both mouse and human (*20–23*). To ensure comparison of similar axial levels in both species, we initially exposed mouse ESCs to a 20h pulse of WNT signalling, and human ESCs to a 72h pulse (*20, 24*). This generated cells with a posterior epiblast identity – so called neuromesodermal progenitors – that express a suite of genes including T/BRA, SOX2 and CDX2 (*20, 25*) (Fig. S1A). These were then exposed to 100 nM of Retinoic Acid (RA), which acts as a neuralizing signal, and to 500nM Smoothened agonist (SAG) that ventralises neural progenitors (*26*)(Fig. 2A,B). For both mouse and human, this resulted in the efficient generation of MN progenitors (pMN) expressing OLIG2 (Fig. 2C, D, S1C), and MNs expressing ISLET1 (ISL1), HB9/MNX1 and neuronal class III beta-tubulin (TUBB3) (Fig. 2E). Progenitors that had not differentiated into neurons switched from Olig2 expressing pMN to p3 progenitors expressing NKX2.2 (Fig. 2C, D). Mouse and human MNs expressed HOXC6, characteristic of forelimb level spinal cord MNs (*27*) (Fig. 2F), indicating pMN and MNs with similar axial levels were being produced in both cases.

**Figure 2.**
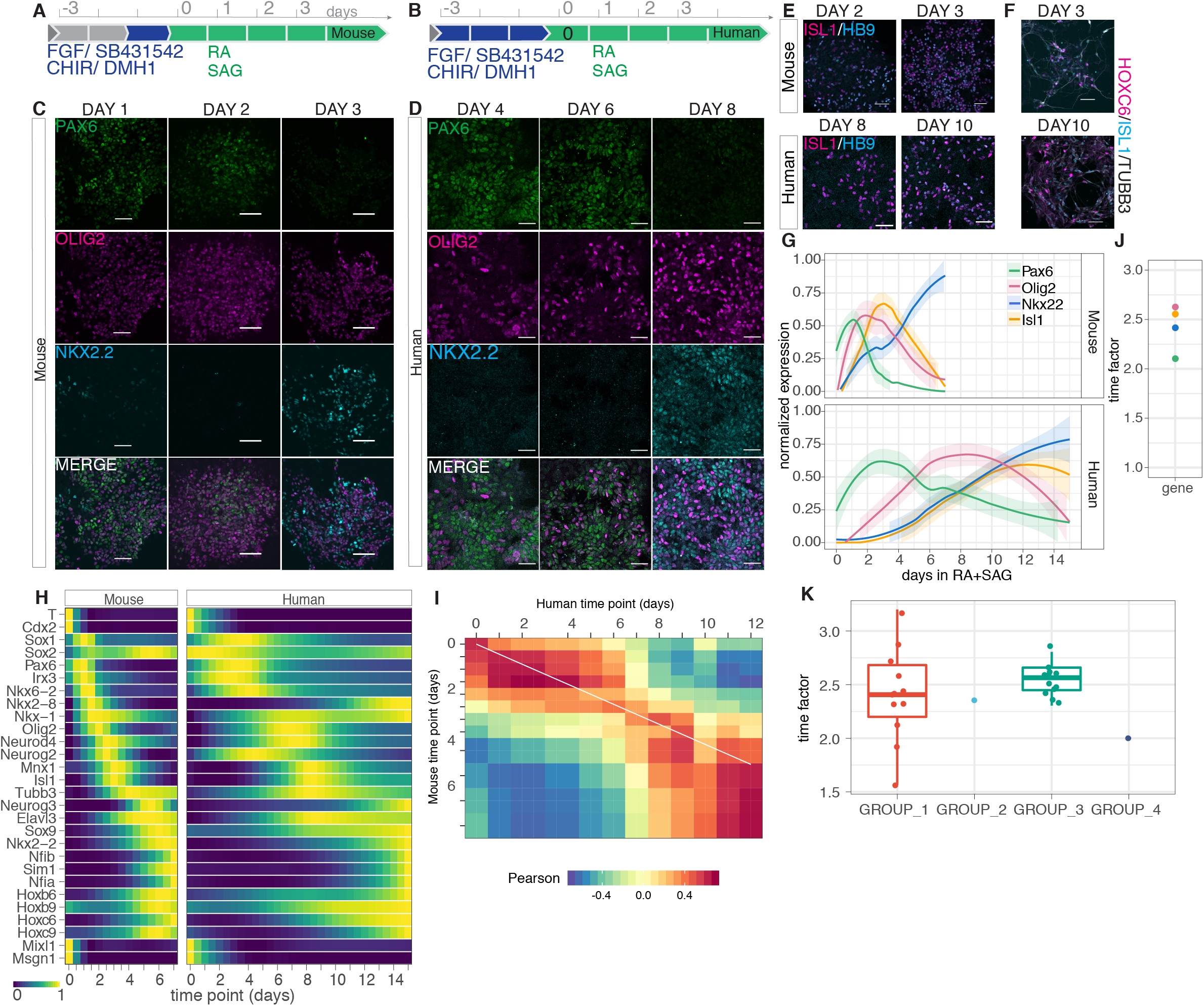
A global scaling factor for in vitro differentiation of mouse and human MNs. (**A**) Schematic of mouse ESCs differentiated to MNs. Spinal cord progenitors are generated via an NMP state induced by the addition of FGF, WNT and dual SMAD inhibition signals for 24h (blue rectangle), and subsequently exposed to the neuralizing signal retinoic acid (RA) and smoothened agonist (SAG) to ventralise the cells (green). (**B**) Schematic of the analogous strategy used for human ESCs to generate MNs, where the addition of FGF, WNT and dual SMAD inhibition signals lasts 72h. (**C**) Expression of NP markers (PAX6, OLIG2, NKX2.2) between Days 1 and 3 in mouse MN differentiations. (**D**) Expression of NP markers (PAX6, OLIG2, NKX2.2) at Days 4, 6 and 8 in human MN differentiations. (**E**) Expression of MN markers (ISL1, HB9/MNX1) in mouse and human MNs. Mouse MNs can be detected by Days 2-3, whereas human MNs are not detected until days 8 and 10. (**F**) HOXC6 expression in MNs characterized by ISL1 and TUBB3 expression at day 3 in mouse and in human day 10. Scale bars = 50 μm. (**G**) RT-qPCR analysis of Pax6, Olig2, Nkx2.2 and Isl1 expression in mouse and human differentiations reveals a conserved progression in gene expression but a different tempo (human n=9, mouse n=9). (**H**) Heatmap of RNA-seq data across mouse and human MN differentiation indicating the interpolated normalized expression of selected markers representative of neuromesodermal progenitors, neural progenitors, neurons, glia and mesoderm cell types. (**I**) Heatmap of pair wise Pearson correlation coefficients of the transcriptomes of mouse (vertical) and human (horizontal) differentiation at the indicated time points. High positive correlation is indicated by values close to 1 (red). White line shows a linear fit of the Pearson correlation over measuring a temporal scaling factor of 2.5 ± 0.2 (median ± std). (**J**) Scaling factor for transcriptome clusters that contain Pax6, Irx3, Olig2, Nkx2.2, Isl1 and Tubb3. (**K**) Time factor estimations for cluster pairs with high proportion of orthologous genes.

Comparison of the two species revealed the same sequence of gene expression changes: expression of Pax6 in newly induced neural progenitors, followed by the expression of the MN progenitor marker Olig2, which precedes the induction of post-mitotic MN markers, including Isl1 (Fig. 2C,G, S1B). But the rate of progression differed. Immunofluorescence and qPCR assays for specific components of the GRN indicated that from the addition of RA and SAG until the onset of ISL1 expression took 2-3 days in mouse but ~6 days in human (Fig. 2E,F,G, consistent with the slower developmental progression in the developing human embryonic spinal cord. Moreover, Olig2 induction peaked at day 2 in mouse but not until day 6-8 in human (Fig. 2G, S1B). Differences in tempo have also been observed between the differentiation of mouse and human pluripotent stem cells (*28*). To test whether the difference in tempo of mouse and human MN differentiation represented a global change in the rate of developmental progression we performed bulk transcriptomics. This revealed a similar pattern of gene expression changes in mouse and human but the changes in mouse cells preceded their human orthologs (Fig. 2H). Cross-species comparison showed a high degree of correlation although altered in time between mouse and human (Fig. 2I, S1D). Moreover, the relative difference in developmental tempo appears constant throughout the differentiation process suggesting a global temporal scaling – developmental allochrony - between mouse and human.

To relate the tempo of mouse and human MN differentiation, we first estimated the global difference in the tempo of gene expression comparing the Pearson correlation coefficients from the transcriptome analysis of both species. This identified a scaling factor of 2.5 (Fig. 2I). Additionally, we clustered gene expression profiles into sets of similarly expressed genes during the time course and we measured the fold difference in the time of appearance of the clusters that contained Pax6, Irx3, Olig2, Nkx2.2, Isl1 and Tubb3 genes. This confirmed that a scaling factor of ~2.5 fit each of the cluster gene expression profiles (Fig. 2J). Similarly, time factor measurements for selected individual genes identified a scaling factor between 2-3 (Fig. S1F,G). To test if the identified time factor could be extended to the whole transcriptome, we selected four cluster pairs that shared a higher proportion of homologous genes than estimated by chance (adjusted p-value < 0.001) (Fig. S1E). A search for a scaling factor that accommodated the difference in the timing of expression in these groups indicated a factor of 2.5 for each of the clusters (Fig. 2K). Together, these results suggest that MN differentiation can be recapitulated in vitro from mouse and human ESCs using similar conditions, resulting in a global 2.5 fold decrease in the rate at which gene expression programmes advance in human compared to mouse.

### Sonic Hedgehog Signalling Kinetics And Sensitivity Do Not Regulate Tempo

Having identified a global scaling factor for the GRN, we set out to investigate the mechanism that sets the timescale. We reasoned that the mechanism was likely to be cell-autonomous since the temporal differences are observed between mouse and human cells grown in the same conditions in vitro, and it has been shown that in vitro differentiated cells transplanted in a host follow their own species-specific dynamics (*29–31*). Since the directed differentiation towards MNs occurs in response to Shh signalling, we hypothesized that the delay in the GRN in human compared to mouse could be a consequence of a reduced response to signalling. To test whether the human GRN could be sped up by higher levels of signalling, we differentiated human progenitors in the presence of increasing concentrations of SAG and in a combination of SAG and Purmorphamine (Pur), another smoothened agonist (Fig. 3A). Single cell measurements of NKX6.1, a GRN transcription factor induced by Shh in ventral progenitors, showed similar proportions and intensity of expression for all levels of signal at equivalent time points (Fig. S2A,B). To test whether the competence of neural progenitors to respond to Shh was delayed in human compared to mouse, we delayed addition of SAG for 24h. A 24h delay in Shh addition resulted in higher initial levels of *IRX3,* as expected, but did not change the time of *NKX6.1* induction relative to the time of SAG addition (Fig. 3C), corroborating the onset of Shh responsiveness is acquired at neural induction in human as in mouse cells.

**Figure 3.**
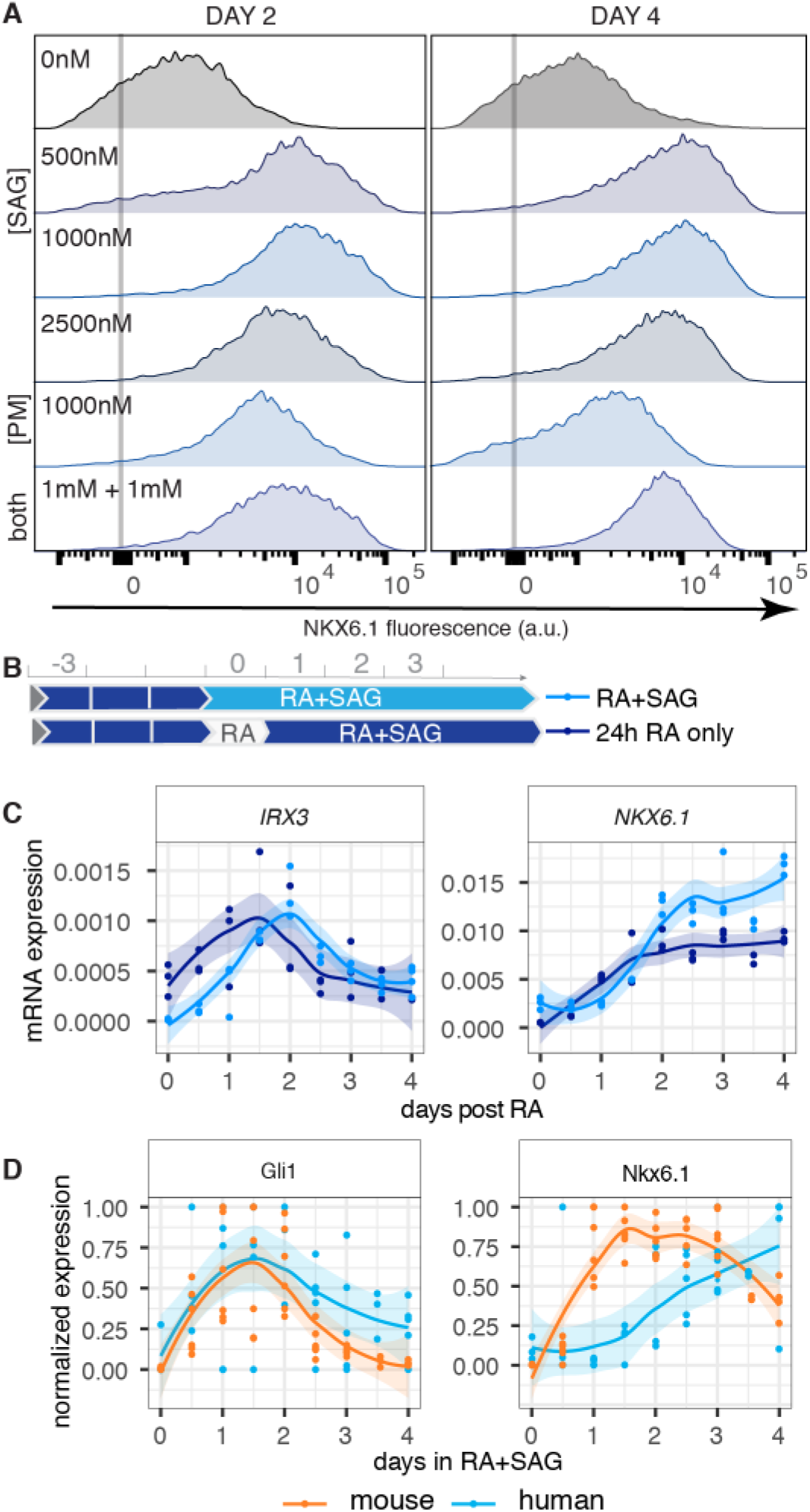
Dynamics of Shh signalling in mouse and human neural progenitors. **(A)** Flow cytometry analysis of NKX6.1 expression in human NPs treated with the smoothened agonists SAG, purmorphamine (PM) or the two combined (both) shows a similar distribution of NKX6.1 expression at days 2 and 4. **(B)** Scheme outlining the standard differentiation protocol, in which RA and SAG are added at the same time (light blue), versus a treatment where SAG addition is delayed for 24h (dark blue). **(C)** RT-qPCR data reveals higher expression of *IRX3* when cells are treated for 24h with only RA (dark blue), whereas there are no substantial differences in the induction dynamics *NKX6.1,* measured from the time of SAG addition. **(D)** RT-qPCR data measured at 12h intervals reveal similar gene expression dynamics in mouse (orange) and human (blue) for Gli1, but distinct for Nkx6.1. (a.u., arbitrary units).

We then compared the kinetics of Shh signalling in mouse and human cells by assaying the response of Ptch1 and Gli1, two Shh pathway components that are Shh direct target genes (*32, 33*). Strikingly, in contrast to Nkx6.1, we found no difference in the response dynamics of these two genes between mouse and human. In both mouse and human cells, the expression levels of Ptch1 and Gli1 were increased within 12h and peaked by 24h (Fig. 3D, S2E). By contrast, the induction of Nkx6.1 was delayed 48h in human compared to mouse (Fig. 3D). Additional Shh targets including Gli2, Ptch2 and Hhip confirmed invariant induction dynamics in mouse and human. The onset of expression for Gli2, Ptch2 and Hhip was detected in less than 24h, and reached maximum expression by 24h (Fig. S2F). By contrast, the induction of Olig2, similarly to Nkx6.1, was delayed in human compared to mouse (Fig. S2F). Together, these results suggest that the onset of intracellular signal transduction does not show species-specific differences and that sensitivity to signals does not appear to have a major role in regulating the tempo of development.

### No Effect Of Interspecies Sequence Differences In Gene Regulation

Having ruled out a role for Shh signalling, we next focused on possible interspecies sequence differences in gene regulation. Even though genes in the GRN are conserved, we hypothesized that sequence differences in the coding region and/or cis-regulatory elements might determine the tempo of development. To study sequence differences between species, we focused our attention on Olig2; it is the major regulator of pMN and regulatory elements for Olig2 have been characterized (*34, 35*). We reasoned that if sequence differences were responsible for the different temporal dynamics in mouse and human cells, we would be able to detect species-specific changes in the timing of Olig2 expression if we introduced the human Olig2 locus into mouse cells. The human Olig2 gene is located on chromosome 21 and we took advantage of the 47-1 mouse ESC line that contains the Hsa21q arm of human chromosome 21 (*36*). We differentiated 47-1 (hereafter referred to as hChr21) alongside its parental line, which lacked Hsa21q, from which it was generated (hereafter referred to as wt). The proportions of neural progenitors and the dynamics of gene expression, measured by RNA expression, immunofluorescence and flow cytometry, were similar between hChr21 and wt lines (Fig. 4A,B,S3A,B). We then assessed the timing of expression of the *hOLIG2* allele. We detected induction of *hOLIG2* at day 1 of differentiation (Fig. 4C), 24h after addition of RA SAG. By contrast in human cells, *hOLIG2* induction is not induced until day 2-3 (Fig. 2G). Thus, in mouse cells, *hOLIG2* follows the same dynamics of gene expression as mouse *Olig2* (*mOlig2*), indicating that the temporal control of gene expression depends on the cellular environment and not the species origin of the genomic sequence.

**Figure 4.**
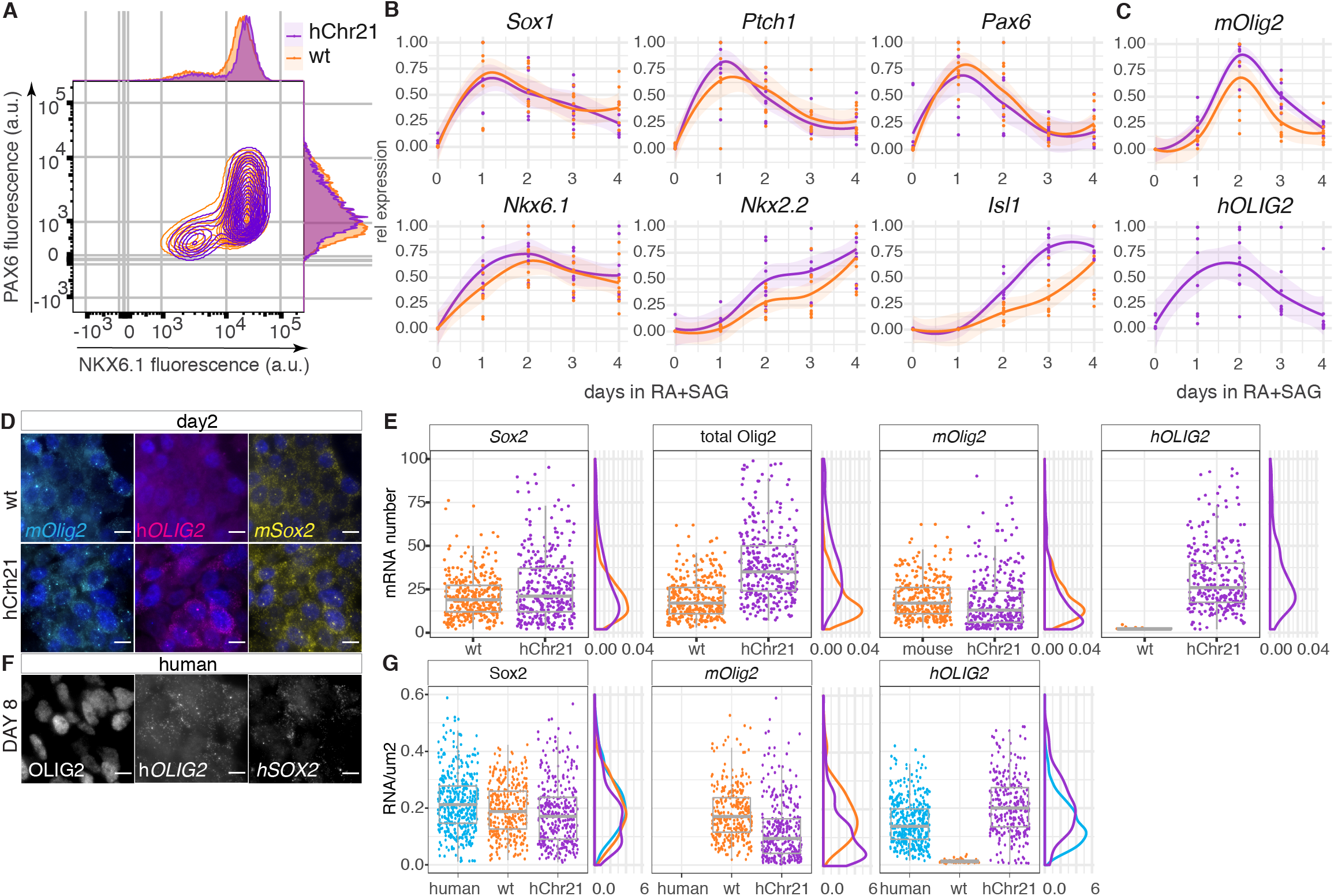
Temporal control of gene expression depends on the species cellular environment. (**A**) Scatter plot with histograms of PAX6 and NKX6.1 expression measured by FACS in neural progenitors from wt (orange) and hChr21 containing (purple) mouse cells at day 2. (**B**) RT-qPCR analysis for the NP and neuronal markers *Pax6, Nkx6.1, Nkx2.2* and *Isl1* in wt and hChr21 lines. (**C**) RT-qPCR expression of Olig2 from the mouse (mOlig2) and human alleles (hOlig2) in wt and hChr21 lines. (**D**) smFISH at day 2 of differentiation in wt and hChr21 lines with probes for *mSox2,* and allele specific detection of *mOlig2* or human Olig2 *(hOLIG2).* Scale bars = 10μm (**E**) Boxplots of RNA molecules per cell in wt and hChr21 cells using Sox2, Olig2 and human- and mouse-allele specific probes. (**F**) smFISH in human NPs at day 8 of differentiation for *hSOX2* and *hOLIG2.* Scale bars = 50μm. (**G**) Boxplots of Sox2 and Olig2 mRNA molecules per cell in human NPs at day 8 and mouse wt and hChr21 cells at day 2.

To compare Olig2 expression levels between the mouse and human alleles, we performed single-molecule Fluorescent In Situ Hybridization (smFISH) (Fig. 4D, S3C). We first assayed transcripts of *Sox2 (mSox2),* a transcription factor expressed in all neural progenitors. The mean and variance in *mSox2* transcripts were similar in both hChr21 and wt neural progenitors, supporting the comparability of the two cell lines (Fig. 4E). We then measured *Olig2* transcripts using species specific probes. The number of mouse *Olig2 (mOlig2)* transcripts in hChr21 cells was lower than in wt cells, but the mean total number of Olig2 transcripts in hChr21 cells, combining mouse and human alleles, was higher than the mean number of transcripts in wt cells (Fig. 4E). This suggests that the number of transcripts that cells express depends on the number of the alleles.

We next asked whether the levels of specific mRNAs were similar in human cells to those in mouse. To this end, we performed smFISH in human neural progenitors for *SOX2 (hSOX2)* and *OLIG2 (hOLIG2)* (Fig. S3D, 4F). The median number of *hOLIG2* molecules in human cells at days 4, 6 and 8 was similar, indicating that the number of transcripts is constant in cells (Fig. S3F). The number of Sox2 and Olig2 transcripts in human neural progenitors were higher than in mouse (Fig. S3E). However, human neural progenitors were larger than mouse progenitors (data not shown) and taking this into account allowed calculation of the concentration of mRNAs (RNAs/μm^2^) in human and mouse cells (Fig. 4G). Strikingly, the median concentration of *hOLIG2* in mouse hChr21 cells was more similar to the concentration of *mOlig2* in wt mouse cells than the concentration of *hOLIG2* in human cells (0.20 in hChr21 cells vs 0.16 in human cells; 0.19 in wt cells), indicating that mRNA might be controlled by the cellular context (Fig. 4G). Overall, we conclude that gene regulation in mouse cells follow mouse-specific characteristics, irrespective of the species origin of the allele, suggesting that the species differences in gene expression dynamics are not encoded within the regulatory genome of individual genes.

### Kinetics Of The Proteome Correspond With The Interspecies Dynamics Of Differentiation

Given that the species difference in tempo did not appear to depend on the specific genomic elements, we reasoned that kinetic features of gene expression must explain the difference. We therefore set out to measure the degradation rate of transcripts and proteins in mouse and human neural progenitors. To assay RNA stability, we used the fluorescent uridine analogue, 5-ethynyluridine (EU) and assayed mouse neural progenitors from day 2 and human neural progenitors from day 4 and 8, representing equivalent developmental states in the two species (Fig. S1C). We pulsed cells for 3h to label actively transcribing mRNAs, transferred them to media lacking EU and collected samples at regular timepoints to assay the level of EU remaining in cells (Fig S4A,B). FACS analysis revealed no differences in global mRNA stability between mouse and human neural progenitors, with a median half-life (t1/2) of 97.75 ± 33.3 min in mouse cells and a t1/2 of 79 ± 19.7 min in human day 4 and 106.45 ± 37.6 min day 8 (Fig 5A,B). Consistent with this, measuring the stability of selected individual mRNAs confirmed the similarity in half-life in mouse and human neural progenitors (Fig. S4E). These quantifications agree with measurements of mRNA half-lives in murine and human immortalized cell lines (*37*).

**Figure 5.**
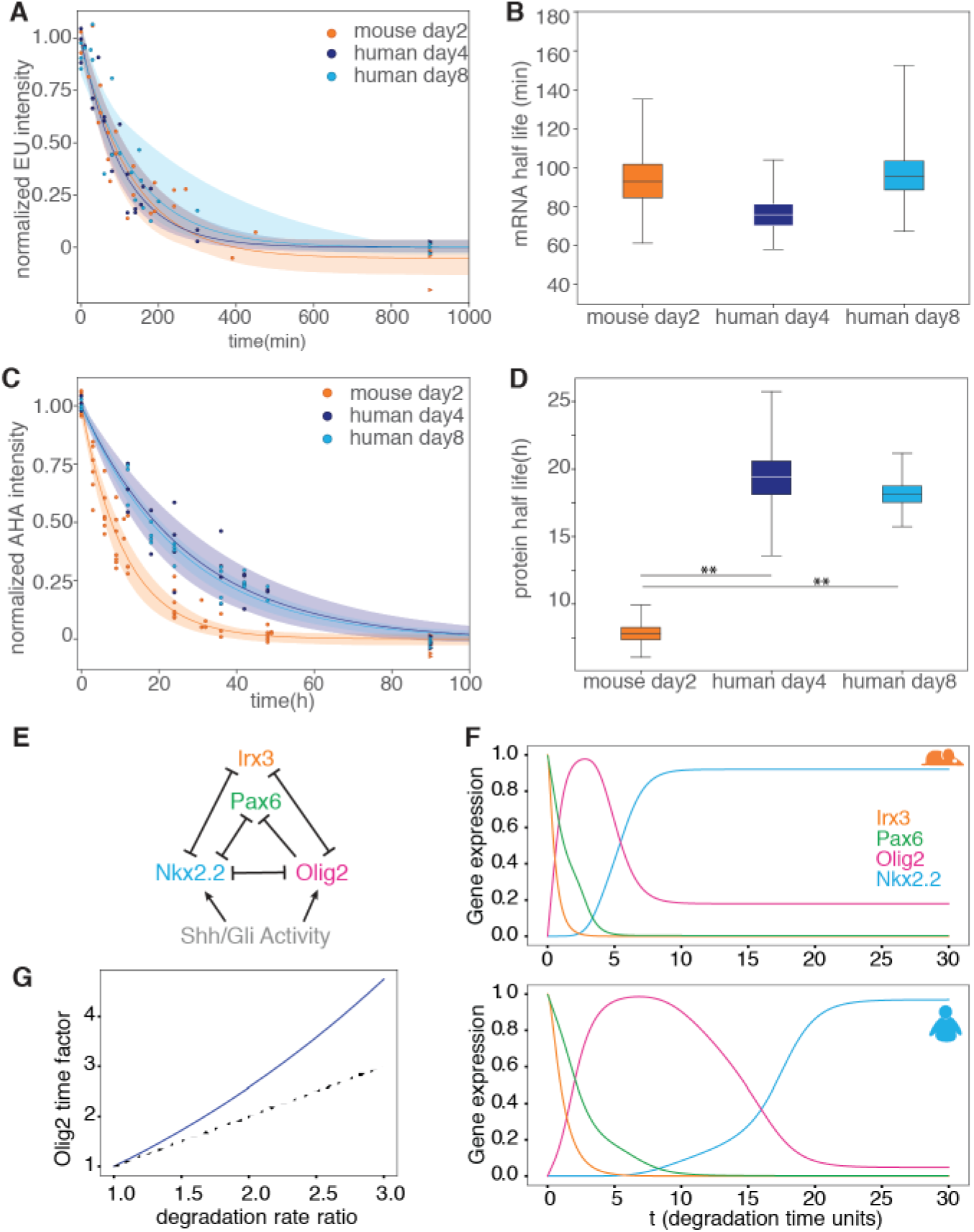
Protein stability in the GRN corresponds to tempo differences between species. (**A**) Normalized EU incorporation intensity measurements to estimate mRNA half-life in mouse (orange) and human (blue) neural progenitors. Line and shadowed areas show best exponential fit and its 70% High Density Interval (HDI). **(B)** Half-life of the transcriptome in mouse neural progenitors at day two (orange), and human neural progenitors at day 4 (dark blue) and day 8 (light blue). (**C**) Normalized AHA intensity measurements of the proteome in mouse (orange) and human (blue) neural progenitors to estimate protein stability. **(D)** Global stability of the proteome in mouse neural progenitors at day two (orange), and human neural progenitors at days 4 (dark blue) and day 8 (light blue). Statistical significance (**) corresponds with <1% of overlap between the distributions of parameter estimations. **(E)** Diagram of the cross-repressive GRN comprising the TFs Pax6, Olig2, Nkx2,2 and Irx3 used to model ventral patterning of the neural tube. **(F)** Temporal dynamics of the simulated GRN model at a relative dorsoventral (DV) position x=0.2 for the mouse model, and the predicted human behaviour by halving the degradation rates of the proteins of the network. **(G)** Estimated time factor at a DV position x=0.2 plotted as a function of the fold change in the degradation rate ratio (blue solid line). The dashed line corresponds to the linear relationship where the degradation rate is the same as the time factor.

Next, we tested whether differences in the half-life of proteins comprising the GRN could explain the allochrony. To assay protein stability of endogenous proteins from mouse and human progenitors, we metabolically labelled nascent proteins replacing methionine in the medium with the methionine analog L-azidohomoalanine (AHA), and used FACS to measure the stability of newly synthesized proteins upon removal of the amino acid analog over the course of 48h (Fig S4C,D). We found that the half-life of the proteome in mouse neural progenitors was shorter than in human progenitors (t_1/2_ = 8 ± 1.6 h in mouse versus t_1/2_=20.5 ± 5.2h in human day 4 or t_1/2_=18.5 ± 2.4 in human day 8), an approximate 2.5 fold difference (Fig. 5C,D). This identifies a global difference in the protein lifetime between mouse and human that corresponds to the difference in tempo.

To test whether changes in protein stability could account for differences in developmental tempo, we took advantage of a mathematical model of the GRN, which we had previously developed (*38*)(Fig. 5E). Doubling the stability of the TFs to mimic human kinetics, resulted in a slower dynamic of the network with the same sequence of gene expression, comparable to that observed experimentally (Fig. 5F). To explore further the connection between changes in protein stability and GRN dynamics we measured the change in time of the onset of Olig2 as a function of degradation rate. This revealed a superlinear relationship in which an increase in protein stability slows GRN dynamics by slightly more than the fold increase in degradation rate (Fig. 5G, S4F). This confirms that an increase in protein stability can explain the tempo changes in MN differentiation between mouse and human. In addition, the analysis revealed that large changes in protein stability can lead to different sequences of gene expression, predicting limits to the allochronies compatible with the GRN (Fig. S4F).

A prediction that arises from this analysis is that the TFs comprising the GRN that regulate MN differentiation should be more stable in human than mouse neural progenitors, and that a fold increase of protein stability close to 2 would give a scaling factor of ~2.5. To this end, we performed pulse-chase experiments labeling nascent proteins with AHA, conjugated labelled proteins to biotin and then streptavidin agarose beads to purify them This revealed that pan-neural proteins SOX1 and SOX2 had longer lifetimes than OLIG2 and NKX6.1 proteins in both species (Fig. 6A,S4G,H). Moreover, human NKX6.1 and OLIG2 were ~2 fold more stable than their mouse homologues (mNKX6.1 ≈2.5 vs. hNKX6.1≈6h; mOLIG2 ≈3.5h, hOLIG2 ≈ 6.8h) (Fig. 6A,S4G,H). These results are consistent with the predictions made by the model, revealing a non-linear relationship between degradation rates and tempo scaling.

**Figure 6.**
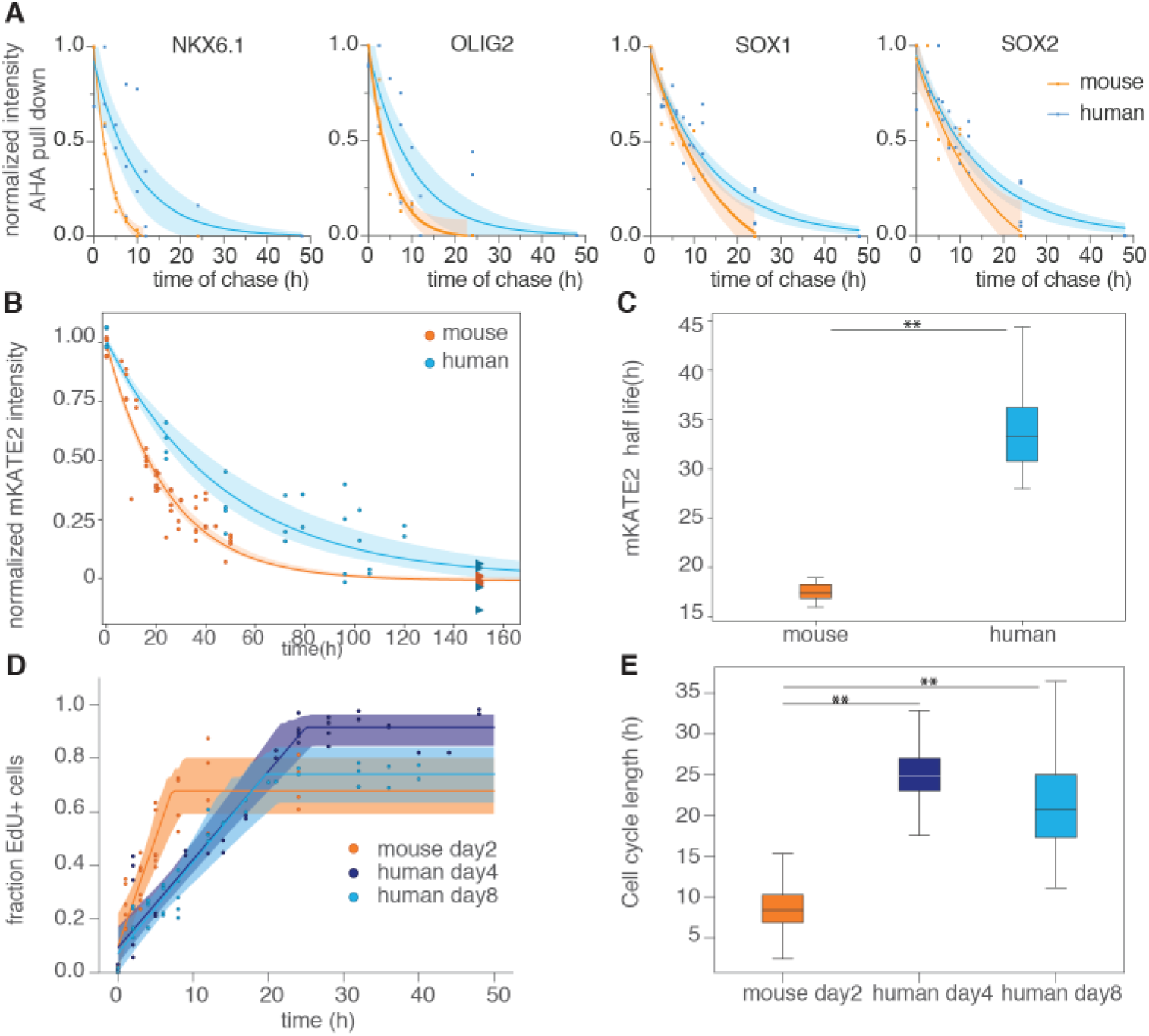
Protein degradation and cell cycle account for the speed differences between species. **(A)** Mouse and human normalized intensity measurements of NKX6.1, OLIG2, SOX1 and SOX2 after AHA pulse-chase experiments on AHA-labeled and purified nascent proteins. Line and shadowed areas show best exponential fit with 95% confidence intervals. **(B)** Normalized intensity measurements of mKATE2 in mouse and human Ptch1::T2A-mKate2 cell lines. Line and shadowed areas show best exponential fit and its 70% HDI. **(C)** Estimated half-lives for mKATE2 in mouse (orange) and human (blue) cells. **(D)** Cell cycle measurements of mouse neural progenitors at day two, and human neural progenitors at days 4 and 8. Line and shadowed areas show best fit and its 80% HDI **(E)** Cell cycle length estimations in mouse neural progenitors at day two, and human neural progenitors at days 4 and 8. For all plots, mouse data is orange-colored, and human is blue. Statistical significance (**) corresponds with <1% of overlap between the distributions of parameter estimations.

The identification of a global increase in the lifetime of proteins in human compared to mouse neural progenitors raised the possibility that exogenous proteins would show species-specific stability. To this end, we generated Patched1-mKate2 reporter lines in mouse and human stem cells. In these lines, we fused the monomeric far-red fluorescent protein Katushka-2 (mKate2) to the C-terminus of endogenous Ptch1 via a self-cleaving peptide (Fig. S5A). This way, we could modulate mKATE2 expression, driven by the Shh responsive Ptch1, using small molecule activators and inhibitors of Sonic Hedgehog signalling. To measure mKATE2 lifetimes, we induced mKate2 expression by addition of SAG (Fig. S5B). Next, we added the Smoothened antagonist Vismodegib (*39*) to block Shh signalling and thereby repress new mKATE2 production. We then assayed the decay of mKATE2 fluorescence in inhibited cells. FACS analysis showed a half-life of 17.8 ± 2.3 h for mKate2 in mouse cells. By contrast, the half-life of the same mKATE2 exogenous protein in human cells was 35.09h ± 7.3 h (Fig 6B,C). These results indicate that protein half-life is species specific.

The long half-life of mKATE2 raised the possibility that dilution, following cell division, contributed to the measured decay rate (*40*). Differences in the cell cycle time between mouse and human neural progenitors could therefore contribute to the difference in mKATE2 lifetime. To test this, we assayed total cell cycle length using cumulative EdU labelling of mouse and human neural progenitors (Fig. 6D,E, S5C,D) (*41*). Cell cycle duration in equivalent staged neural progenitors from mouse and human was 10.8h ± 8.3h compared to 28.4h ± 13.9h, respectively. Thus, similar to the proteome, the cell cycle operates twice as fast in mouse compared to human. Since progress through the cell cycle is controlled by protein degradation (*42, 43*), the difference in cell cycle rate between mouse and human cells may also be a consequence of a global change in protein stability.

Taken together, these data indicate that the dynamics of the gene regulatory network associated with the embryonic generation of MNs progresses 2-3 times faster in mouse than in human cells. A similar difference in the tempo of the segmentation clock between mouse and human has also been observed (*9, 11*). These differences do not appear to arise from a bottleneck caused by a specific rate limiting event in MN generation. Moreover, neither changes in the dynamics of signalling nor variations in genomic regulatory sequences appear to account for the species-specific tempos. Instead, the correlated ~2.5 fold differences in cell cycle length and general protein stability suggest that the temporal scaling in developmental processes results from global differences in key kinetic parameters that broadly affect the tempo of molecular processes. What sets this global tempo remains to be determined but could involve the differences in the rates of pivotal molecular processes such as global changes in proteostasis or differences in the overall metabolic rate of cells. How these affect the pace at which GRNs elaborate and how such variations are assimilated to ensure the development of robust and appropriately proportioned tissues will need to be addressed, but the availability of in vitro systems that mimic in vivo developmental allochrony open up the possibility of exploring these issues.

## ACKNOWLEDGEMENTS

We are grateful for the human embryonic material provided by MRC/Wellcome Trust (MR/R006237/1) Human Developmental Biology Resource and the generous donors whose contributions have enabled part of this research. We thank Andrés de la Peña, Anestis Tsakiridis, Vicki Metzis, Andreas Sagner, Tom Watson, M. Joaquina Delás, Robert Blassberg, Julien Delile, Tristan A Rodríguez, Phillip East, and Robert Goldstone as well as other members of the lab for advice, reagents and critical feedback. We thank the Crick Science Technology Platforms in particular the Advanced Sequencing Facility, the Equipment Park, the Flow Cytometry Facility, and the Bioinformatics and Biostatistics group.

## AUTHOR CONTRIBUTIONS

T.R. and J.B. conceived the project, interpreted the data, and wrote the manuscript with input from all authors. T.R. designed and performed experiments and data analysis. D.S. designed and performed experiments and data analysis. R.P.C. performed theoretical modelling and data analysis. L.G.P. designed experiments and performed data analysis for smFISH. C.B. performed bioinformatic analysis. M.M. performed embryo work, generated and characterized the Ptch1::T2A-mKate2 mouse ES cell line. K.E. performed embryo work. E.M. and V.T. provided reagents and feedback.

## DATA AND MATERIALS AVAILABILITY

The accession number for the bulk RNA-seq data reported in this paper is GSE140749.

## SUPPLEMENTARY MATERIALS

Materials and Methods

Fig S1 - S5

Table S1 - S2

### METHODS

#### Tissue Preparation

Human embryonic material (4-6 weeks of gestation) was obtained from the MRC/Wellcome-Trust (grant #006237/1) funded Human Developmental Biology Resource (HDBR57, http://www.hdbr.org) with appropriate maternal written consent and approval from the London Fulham Research Ethics Committee (18/LO/0822) and the Newcastle and North Tyneside NHS Health Authority Joint Ethics Committee (08/H0906/21+5). HDBR is regulated by the UK Human Tissue Authority (HTA; www.hta.gov.uk) and operates in accordance with the relevant HTA Codes of Practice. Tissue was fixed in 4% paraformaldehyde (PFA) overnight at 4°C, washed in PBS, and transferred to 15% sucrose in phosphate buffer overnight at 4°C. Embryos were subsequently embedded in gelatin solution (7.5% gelatin, 15% sucrose in phosphate buffer) and snap-frozen in isopentane on dry ice. Transverse cryosections (thickness: 14μm) were cut using a Leica CM3050S cryostat (Leica Microsystems Limited, Milton Keynes, UK) and placed on Superfrost Plus™ slides (Thermo Scientific™ 10149870). Slides were stored at −80°C until ready to be processed for immunohistochemistry. Following immunohistochemistry 22 x 50 mm No.1,5 thickness coverslips (VWR 631-0138) were mounted on to the sections using ProLong™ Gold antifade reagent (Invitrogen P36930).

Mouse spinal cord tissue was prepared in the same way as described. All animal procedures were carried out in accordance with the Animal (Scientific Procedures) Act 1986 under the Home Office project license PPL80/2528 and PD415DD17.

#### Immunostaining and microscopy

Immunohistochemistry on human and mouse spinal cord tissues, and on mouse and human cells was performed as described previously (*44*). Primary antibodies were diluted as follows: rabbit anti-Pax6 (Covance 1:500), goat-anti Olig2 (R&D AF2418, 1:800), mouse anti-Nkx2.2 (,1:500), mouse anti-Hb9/Mnx1 (DSHB, 1:40), goat anti-Isl1 (R&D AF1837, 1:1,000), mouse anti-Hoxc6 (Santa Cruz Biotechnology sc-376330, 1:200), mouse anti-Tubb3 (Covance MMS-435P, 1:500), chicken anti-Tubb3 (Abcam ab107216, 1:500), goat anti-sox2 (R&D, 1:500), goat anti-sox9 (R&D AF3075, 1:250), rabbit anti-NFIA (Atlas antibodies HPA008884, 1:500). Alexa secondary antibodies (Life Technologies or Jackson Immunoresearch) used throughout this study were diluted 1:1,000, and nuclei were counterstained with DAPI.

Cryosections were imaged using a Leica SP8 confocal microscope equipped with a 20x NA 0.75 dry objective, or a Leica SP5 confocal microscope. Cells were imaged using a Zeiss Imager.Z2 microscope using the ApoTome.2 structured illumination platform or using a Leica SP5 confocal microscope. Z stacks were acquired and represented as maximum intensity projections using ImageJ software. Pixel intensities were adjusted across the entire image in Fiji. The same settings were applied to all images. Immunofluorescence was performed on a minimum of 3 biological replicates, from independent experiments.

#### Cell culture and neural progenitor differentiation

All mouse ESCs (HM1, D3 and Ptch1::T2A-mKate2 mouse ESC line) were propagated on mitotically inactivated mouse embryonic fibroblasts (feeders) in DMEM knockout medium supplemented with 2000U/ml ESGRO-LIF (ESG117 Sigma Aldrich), 10% cell-culture validated fetal bovine serum, penicillin/streptomycin, 2mM L-glutamine (GIBCO). The 47-1 (hChr21) cell line (*36*) was maintained with 0.8ul/mL of G418 (GIBCO) to select for cells carrying the human chromosome 21.

H9 ESC line (WiCell), and Ptch1::T2A-mKate2 H9 ESC lines were routinely cultured in Essential 8™ medium on 0.5ug/cm2 laminin-coated plates (Thermo Fisher A29249), and split using Versene (Gibco 15040066).

To obtain mouse neural progenitors of posterior identity, mouse ESCs were dissociated with 0.05% trypsin, feeders removed by differential binding to gelatin-coated plates, and 60,000-80,000 cells were plated per 35mm CELLBIND dish in N2B27 medium supplemented with 10ng/ml of bFGF (100-18B Peprotech) from days –3 to –1. On day −1, N2B27 was supplemented with 10ng/ml bFGF (100-18B Peprotech), 5μM CHIR99021 (Axon), 10uM SB431542 (LT S0400) and 2uM DMH1 (A12820 Adooq Bioscience). From day 0 onwards cells and cultured in N2B27 supplemented with 100nM RA (R2625 Sigma Aldrich) and 500nM SAG (566660 Calbiochem) for neural induction. Medium was changed daily. N2B27 medium contained a 1:1 ratio of DMEM/F12:Neurobasal medium (21331020 and 21103049 ThermoFisher Scientific) supplemented with 1xN2 (17502001 ThermoFisher Scientific), 1xB27 (17504001 ThermoFisher Scientific), 2mM L-glutamine (25030024 ThermoFisher Scientific), 40mg/ml BSA (A7979 Sigma Aldrich), Penicillin/Streptomycin (15140122 ThermoFisher Scientific) and 0.1mM 2-mercaptoethanol (21985023 ThermoFisher Scientific).

For human motor neuron differentiation, cells were split with Versene and plated as clusters of ~20 cells on 1ug/cm2 fibronectin-coated (Millipore FC010) wells in N2B27 medium supplemented with 3 μM CHIR99021, 5ng/ml bFGF (100-18B Peprotech), 10uM SB431542 (LT S0400), 2uM DMH1 (A12820 Adooq Bioscience), and 10 μM ROCK inhibitor (Y-27632 Tocris) from days −3 to −1. Rock inhibitor was removed after 48h. On day 0 the cells were dissociated using accutase (A1110501 ThermoFisher Scientific) and 175,000 cells were plated per fibronectin-coated 35mm dish on N2B27 medium containing 100nM RA (R2625 Sigma Aldrich) and 500nM SAG (566660 Calbiochem) with10 μM ROCK inhibitor (Y-27632 Tocris). Rock inhibitor was removed 48h after plating and medium was replaced every other day. All experiments involving hES cells have been approved by the UK Stem Cell Bank steering committee (ref SCSC14-18).

#### Generation of Ptch1::T2A-mKate2 mouse and human ESC lines by CRISPR

For CRISPR/Cas9-mediated homologous recombination, short guide RNA (sgRNA) sequences (mouse: GTGGGGGAGCAGCTCCAACTG, human: GTGAGTGCCACTGACAA) was cloned into pSpCas9(BB)-2A-Puro (Addgene pX459 plasmid # 62988). As donor vector, the T2A-3xNLS-FLAG-mKate2 cassette (*26*) was inserted at the 3’ end of the Patched1 open reading frame, using 2.1 Kbs upstream and 2.8 Kbs downstream arms for mouse and 1.88 Kb upstream and 0.98 Kb downstream homology arms for human.

For mouse ESC targeting, HM1 ESCs were electroporated as in Gouti et al 2017 (*45*). For human ESC targeting, 2×10^6^ cells were electroporated with 2μg of each plasmid using program A23 of Nucleofector II (Amaxa) and Human Stem cell nucleofector I kit (VPH-5012 Lonza). Electroporated cells were plated on matrigel coated 6-well plate and maintained in mTESR and 10 μM ROCK inhibitor (Y-27632 Tocris). For selection, colonies were first treated with 0.5 μg/ml Puromycin (Sigma) for two days followed by 50 μg/ml Geneticin (Gibco) selection. Then the colonies were dissociated using accutase and plated at low density to allow for clonal growth. Individual colonies were picked using a 20-μl pipette tip, dissociated in accutase (Gibco), and replated to allow expansion and a second round of Geneticin selection. Correct integration of the T2A-3xNLS-FLAG-mKate2 transgene was verified using long-range PCRs and Sanger sequencing.

#### RNA extraction, cDNA synthesis and qPCR analysis

RNA was extracted from cells using a QIAGEN RNeasy kit, following the manufacturer’s instructions. Extracts were digested with DNase I to eliminate genomic DNA. First strand cDNA synthesis was performed using Superscript III (Invitrogen) using random hexamers and was amplified using PowerUp SYBR green (Applied Biosystems). qPCR was performed using the QuantStudio 5 Real-Time PCR System. For mouse samples, expression values for each gene were normalized against b-actin, using the delta-delta CT method. For human samples, expression values for each gene were normalized against b-Actin and Gapdh. For human and mouse comparisons, data was normalized to its maximum levels. Error bars represent standard deviation across at least three biological replicate samples in triplicates. Mouse and human qPCR primers used are listed in TableS1. Data were processed, normalized and plotted using R.

#### RNA-seq library preparation and data analysis

Total RNA was purified using the RNeasy Plus Micro Kit (QIAGEN) according to the manufacturer’s instructions. Three separate RNA libraries (biological replicates) were barcoded and prepared for each time point. Indexed libraries were pooled and sequenced on an Illumina HiSeq 4000 flow cell conFig.d to generate 101 cycles of single-ended data. Raw data was demultiplexed and FastQ files created using bcl2fastq (2.20.0). Technical replicates between lanes on the flow cells were merged by concatenation of the FastQ files prior to analysis into biological replicate datasets, each of which contained 30-45 million reads.

Biological replicate datasets were analysed using the BABS-RNASeq (Bah et al., https://github.com/crickbabs/BABS-RNASeq) Nextflow (*46*) pipeline developed at the Francis Crick Institute. The GRCm38 mouse and the GRCh38 human reference genomes were used with the Ensembl release-89 (*47*) gene annotations. Dataset quality was assessed by FastQC (0.11.7, Andrews, http://bioinformatics.babraham.ac.uk/projects/fastqc), RSeQC (2.6.4, *38*), RNA-SeQC and Picard (2.10.1, http://broadinstitute.github.io/picard) and expression was quantified by STAR (2.5.2a, *40*) and RSEM (1.3.0, *41*) using the BABS-RNASeq pipeline.

DP_GP_cluster (*52*) was used to cluster the mean estimated TPM across time courses using default parameters and the --fast, --true_times and --check_convergence options. Genes other than Isl1, Nkx2-2, Olig2 and Pax6, whose range of TPM values was 50 or more, were considered for this analysis. Enrichment of homologous genes between expression clusters was assessed using the phyper() function in R. After multiple testing correction by FDR a significance threshold of 0.001 was applied to identify homologous pairs. Groups of connected homologous pairs were identified using cluster() in igraph (1.2.2, *43*) for R.

##### Time factor estimations

The Pearson correlation time factor (*f*) was obtained by minimizing the Euclidian distance between the linear relationship *t*_ and each inter-species time point pair weighted with the Pearson correlation coefficient of each pair. Error was calculated by bootstraping.

Time scaling factors *f* between RNA-seq profiles of genes and clusters were obtained from the normalized gene expression temporal profiles *y*(*t*), by minimizing the distance *d*(*y_human_,y_mouse_*) between the human expression profile and the scaled profile of mouse *d* = ∫ (*y_human_*(*t*) - *A y_mouse_*(*t · f + B*))^2^ d*t*. Continuous profiles were obtained using Akima spline interpolation, while the minimization of the distance was performed using the Nelder-Mead method. The reported ensemble of time scaling factors correspond with each possible inter-species pair of replicates.

#### Single molecule RNA FISH

Mouse NPs were seeded on Matrigel-coated (GELTREX) coverslips, and human NPs were plated on fibronectin-coated coverslips. Samples were washed in 1 mL of 1X PBS and the coverslip was fixed in 1 mL of Fixation Buffer (4% paraformaldehyde in 1X PBS) at room temperature for 15 min. The fixed cells were washed with 1X PBS and then permeabilized at 4°C for at least 24h using 70% (vol/vol) ethanol. Cells were subsequently rehydrated with 1 ml of Wash Buffer (10% formamide, 2xSSC and 0.25% Triton in water) for 15 min before incubating in 100 μL of Hybridization Buffer (nuclease free water + 10% dextran sulfate + 10% formamide + 2xSSC buffer) containing 125 nM probe and goat-anti Olig2 (R&D AF2418, 1/800) at 37°C in the dark overnight. The following day the coverslips were washed for 30 min at 37°C in wash buffer with Alexa Fluor 488 secondary antibody (1/2000). Nuclei were counterstained with DAPI for 5 min. Coverslips were washed once with GLOX solution (Tris pH 8.0 10 mM, 2xSSC buffer, 0.4% glucose) and mounted on Antifade solution (GLOW with 1μl of Catalase (Sigma-Aldrich, C3515) and 1μl of Glucose Oxidase(Sigma-Aldrich, G2133; dissolved in 50mM sodium acetate, pH 5.5, to a concentration of 3.7 mg/mL)). Mouse-specific and human-specific Olig2 and Sox2 probes conjugated to Quasar 570, CAL Fluor Red 610 or Quasar 670 were designed using the Stellaris Probe Designer online tool and ordered from LGC Biosearch Technologies. A list of the individual probes in the probe sets used for this study is available in TableS2.

Samples were imaged with an Olympus widefield BX61 microscope equipped with a 60x oil UPlanFL N objective (with a 0.9 numerical aperture), an iXon Ultra EMCCD camera (Andor Technology) and a BX2 filter cube with the following filters specific for smFISH: 49304 (Gold FISH), 49310 (Red-2 FISH) and 49307 (Far Red FISH), all from Chroma Technology Corporation. 8.1 μm z-stacks were acquired, with a 0.3 μm distance between individual optical planes, for each wavelength, starting with DAPI, Alexa Fluor 488, Quasar 670, then CAL Fluor Red 610, and then Quasar 570. Typical illumination times are 1.5 s for Quasar 670, 1 s for CAL Fluor Red 610, 1 s for Quasar 570, 200 ms for Alexa Fluor 488 and 1 ms for DAPI. The acquired planes for each Z-stack were focused to ensure maximum coverage of cellular volume and minimum empty areas. The microscope, camera and hardware were controlled through the Micro-Manager software (ImageJ).

A minimum of 10 Z-stacks were acquired per dataset. For quantifications, OLIG2+ cells were manually segmented with Icy (*54*) on maximum intensity projections of the images. The output of Icy plugin consisted of individual.xls files for each location and fluorescence channel imaged. Each file contained the spot XY coordinates, spot mean fluorescence intensity and spot area for all the spots detected within the segmented cells in a given image. A custom-made Python script was used to gather all the information across multiple.xls files in just one file per experiment. The total number of spots detected per cell, which are presumed to be representative of the total number of mRNAs per cell, was obtained by counting the total number of detected spots within each cell, which were uniquely identified in basis of their cell line, sample day, image and cell identifier number. Data analysis and visualization were carried out in R.

#### Intracellular Flow Cytometry

Cells were dissociated with 0.5ml accutase (GIBCO) and fixed in 4% paraformaldehyde in PBS for 5min. For stainings, 1×10^6^ cells were used. Cells were incubated overnight with an antibody mix on PBST with 1% BSA at 4°C. The following day, and when OLIG2 (R&D AF2418, 1:800) primary antibody was used, cells were pelleted and incubated with Alexa Fluor secondary antibodies (1/1000) at room temperature for 1h. Cells were resuspended in 0.5mL PBS and filtered for data acquisition on LSR Fortessa or LSRII cytometers. 10,000 events gated on SOX2 were recorded. The following fluorophore-conjugated antibodies were used: SOX2-V450 (BD 561610, 1:100) PAX6-488 (BD 561664, 1:50), NKX6.1-PE (BD 563338, 1:50). Analysis was performed using FlowJo software. Cells were gated on SOX2 before plotting.

#### Global RNA and protein stability measurements

To measure RNA stability, cells were cultured for 3h in differentiation medium supplemented with 1mM EU. After the pulse, cells were washed once with PBS and cells were grown on fresh medium over the course of the experiment. For proteome stability measurements, cells were starved by replacing complete differentiation medium with methionine-free medium for 30 min. Next, 100μM AHA was added to the methionine-free medium for 1h. To measure protein stability, AHA pulse was removed by washing the cells once with PBS and growing the cells on complete differentiation medium for the course of the experiment. On the indicated days of differentiation and at specific time points after EU or AHA removal, cells were processed for intracellular flow cytometry. EU-incorporated RNAs were labelled using Click-iT™ RNA Alexa Fluor 488 HCS Assay (Thermo Fisher Scientific C10327), and AHA-incorporated proteins were labeled using Click-iT™ Cell Reaction Buffer Kit (Thermo Fisher Scientific C10269) on a volume of 150ul per sample reaction. Estimations of global transcriptome and proteome stability were obtained by normalizing each individual replicate using an initial exponential fit *y*(*t*) = *B* + *C* · exp(*kt*) to determine the baseline *B* and initial fluorescence intensity *C*, allowing a comparison of all the replicates together. Exponential fits of the bootstraps of the normalized ensemble were used to construct credibility distributions of the degradation rates. Error intervals reported correspond with 99% High Density Interval (HDI). A minimum of 2 biological replicates per species per time point from independent experiments were used.

#### Metabolic labeling, biotin conjugation and streptavidin pull down of nascent RNA

Mouse day 2 and human day 8 neural progenitors were pulsed with 1mM of EU for 3h and chased with complete medium for 0 min, 50 min, 90 min and 180 min post pulse. Cells were dissociated using accutase, rinsed in PBS, and pellets resuspended in 600μl of RLT buffer (QIAGEN RNeasy kit) and stored at −80°C until needed. RNA was extracted following the RNA extraction procedure indicated above. Next, the Click-iT Nascent RNA Capture Kit (Invitrogen, C10365) was used to biotinylate and streptavidin purify nascent transcripts. Briefly, 2μg of RNA was clicked to biotin and cleaned-up on a 50<u>μl</u> reaction according to manufacturer’s instructions. For biotinylated RNA binding to streptavidin T1 magnetic Dynabeads, 1ug of RNA was used and 24ul of Dynabeads were used per sample. RNA bound to the beads was immediately used for cDNA synthesis using the SuperScript VILO cDNA synthesus kit (Cat.no. 11754-050) in a 50ul reaction, and cDNA was eluted from the beads by at 85 °C for 5 min. Mouse cDNA samples were diluted 1/5, whereas human cDNA samples were diluted 1/4 for qPCR analysis. For RNA enrichment analysis, expression values for each gene were normalized against ß-actin, using the delta-delta CT method. To compare between biological replicates and species, maximum levels were normalized to 1, and non-linear one-phase decay with 95% confidence intervals curves were fitted using GraphPad Prism version 8. At least two biological replicates from independent experiments were used.

#### Metabolic labeling, biotin conjugation and streptavidin pull down of nascent proteins

Following 1h incubation in methionine free medium, we labeled nascent proteins for 2h with 1mM AHA (L-azidohomoalanine). Next, AHA was washed away and cells either collected for protein extracts or fed with complete differentiation medium then collected for protein extracts at the indicated timepoints. The cells were dissociated using accutase, rinsed in PBS, pellets frozen on dry ice and stored at −80°C until needed. To prepare protein extracts we lysed the cells in 50mM Tris pH8, 1% SDS, 250U/ml benzonase and tablet per 3.5ml of lysis buffer protease inhibitors (cat no 04693159001, Roche). Protein content of the cell lysates was measured using the Pierce BCA protein assay kit (cat no 23225). 1mg of total protein per sample was labeled using biotin alkyne (cat no B10185, Invitrogen) and Click-it technology (Click it Protein Reaction Buffer kit, cat no 10276, Invitrogen). Labelled protein extracts were purified by desalting columns (Zeba 7kDa MWCO, Cat no 89892, ThermoFisher Scientific) and mixed with washed magnetic streptavidin beads (400ul slurry per sample, cat no 65602, ThermoFisher Scientific) for 16h at 4°C. The beads were washed four times with 0.1%SDS, 0.1% BSA PBS, once with 0.1%SDS, 0.1% BSA PBS and biotinylated proteins recovered by heating at 92°C for 10 min in 2x Laemmli buffer.

#### Western Blots and band intensity measurements

Samples were loaded on 12% precast gels (10 wells 4561043 or 15 wells 4561046, Biorad) and run at 110V for 1h 45min. Proteins were transferred on nitrocellulose membrane (cat no 10600012, Amersham) at 300mA for 2h. Membranes were treated with TBS blocking buffer at room temperature for 30min. Primary antibodies were applied in 0.1% Tween-20, TBS blocking buffer for 16h at 4°C. The primary antibodies used were: rabbit Olig2 (Millipore, AB9610, 1/2000), goat Olig2 (R&D, AF2418, 1/1000), mouse Nkx6.1 (DSHB, 1/100), rabbit Nkx6.1 (Novus Biologicals, NBP1-49672, 1/1000), goat Sox1 (R&D Systems, AF3369, 1/500), rabbit Sox2 (Millipore, AB5603, 1/500). Secondary antibodies were applied in 0.2% Tween-20, 0.01%SDS, TBS blocking buffer for 2h at room temperature. The secondary antibodies used (Licor) were: goat anti rabbit 800CW (926-32211, 1/5000), donkey anti mouse 680LT (925-68022, 1/10000), donkey anti goat 800CW (925-32214, 1/5000), donkey anti rabbit 680LT (925-68023, 1/10000). The membranes were stripped when required using NewBlot IR stripping buffer (928-40028, Licor), 30min at room temperature. The membranes were scanned using an Odyssey CLx gel documentation set up. Band intensities were measured using Fiji and plotted using GraphPad Prism version 8 following the procedure described for nascent RNA pull downs. A minimum of 3 biological replicates per species from independent experiments were used.

#### mKATE2 stability measurements

To measure mKATE2 stability, cells were exposed to 100nM RA and 1uM of SAG from day 0 on the neural differentiation protocol to induce maximum levels of Ptch1::T2A-mKate2 expression driven from the Ptch1 locus. Next, Vismodegib (5uM, GDC-0449 Cayman Chemical) was added when mKate2 intensity was maximal, and samples were collected at the indicated time points. Samples were stained and fixed with live/dead cell stain as per manufacturer’s instructions (Life Technologies). mKATE2 intensity was quantified by FACS using the 610/20 yellow red excitation laser on LSR Fortessa cytometers. Analysis was performed using Flowjo software. mKATE2 half-life estimations were performed following the same procedure as for the proteome stability. A minimum of 2 biological replicates in duplicates per species per time point from independent experiments were used.

#### Cell cycle measurements

0.5uM EdU (Thermo Fisher Scientific C10633) was added to mouse and human neural progenitors at the indicated days of differentiation. Samples were then collected in duplicates at the indicated time points, and processed for intracellular flow cytometry. For EdU detection, the Click-iT™ Plus EdU Alexa Fluor™ 488 Flow Cytometry Assay Kit (Thermo Fisher Scientific C10633) was used according to the manufacturer’s instructions, on a volume of 200ul per sample reaction. For EdU incorporation measurements, cells were counterstained with DRAQ5 (Cell Signalling 4084S, 1/10,000) to measure DNA content. Cells were gated on SOX2 and EdU and the percentage of positive cells per time point. Estimates of cell cycle parameters were obtained by assuming that independent cells in the population advance at the same speed along the cell cycle so the fraction (*r*(*t*)) of Edu+ positive cells increases linearly in time until a time equal to the cell cycle duration (T) at which all the responsive cells are stained (*41*) *r*(*t*) = *r_M_* min(1, *r*_0_ + *t*(l – *r*_0_)/*T*). The initial fraction of cells stained at the pulse (*r*_0_), and the maximum fraction of cells labelled in the population (*r_M_*), are parameters of the model. Flat prior distributions for the 3 parameters (*T,r*_0_,*r_M_*), and a Gaussian distributed likelihood around the model prediction, was used to determine the posterior probability distribution of the parameters using PyDream Markov chain Monte Carlo implementation (*55*). The reported parameters in the manuscript correspond with the 99% HDI of the marginal posterior distribution for the cell cycle duration *T*. A minimum of 2 biological replicates in duplicates per species per time point from independent experiments was used.

**Figure S1.**
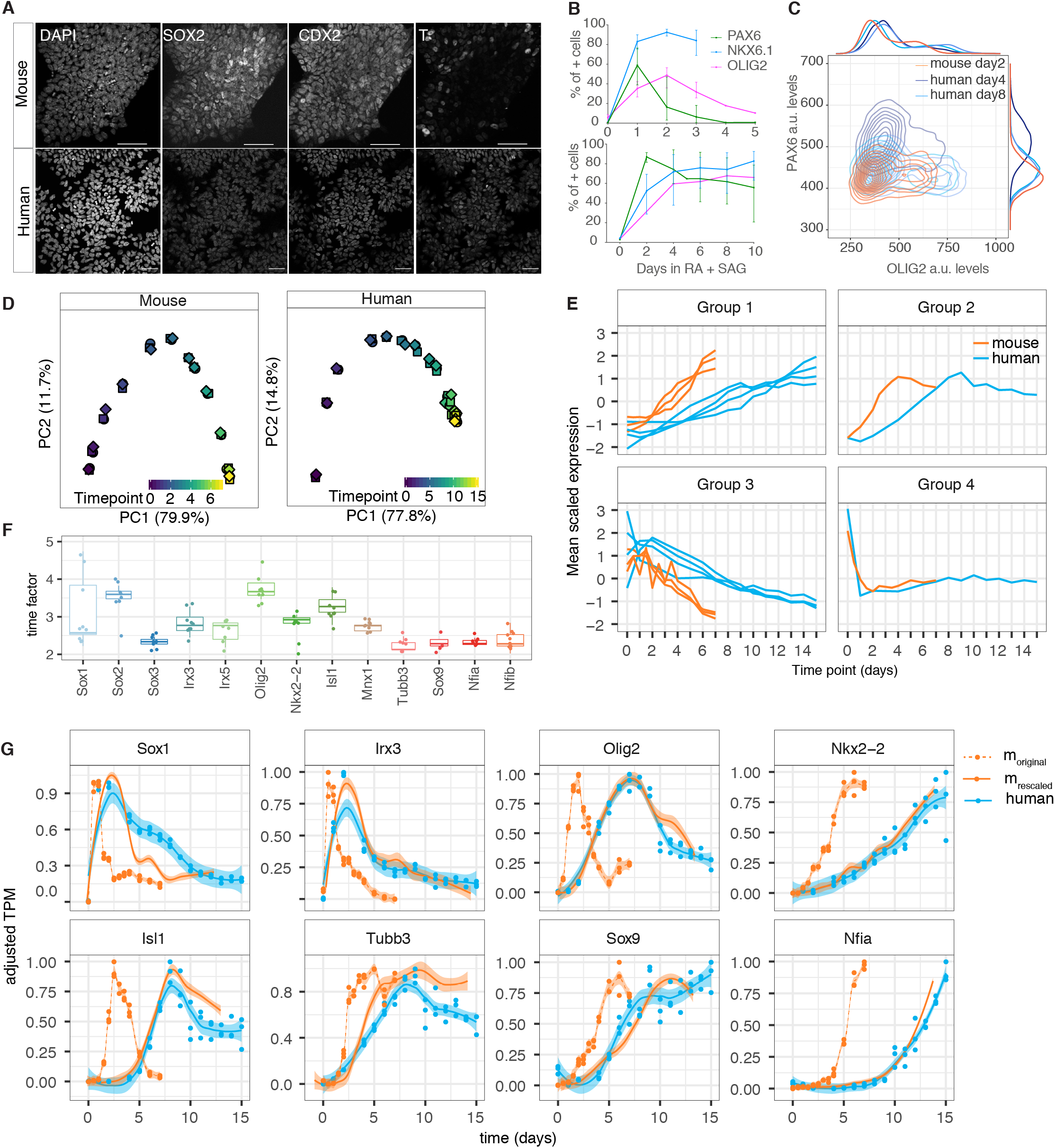
Characterization of a scaling factor for in vitro differentiations from mouse and human cells. **(A)** Expression of the neuromesodermal progenitor markers SOX2, CDX2, T at day 0 in mouse and human motor neuron differentiations. **(B)** Percentage of PAX6, NKX6.1 and OLIG2 positive cells at various time points across mouse and human differentiations. **(C)** Scatter plot with histograms of PAX6 and OLIG2 intensity levels measured by FACS in NPs from mouse cells at day 2 (orange) and human cells at days 4 (dark blue) and day 8 (light blue). **(D)** PCA plots of RNA-seq data across mouse and human MN differentiation. Shapes of the points indicate biological replicates. **(E)** Mean scaled expression profiles of selected human and mouse clusters pairs selected that share a higher proportion of homologous genes than expected by chance. **(F)** Estimated time factor on selected genes measured from the RNAseq dataset. **(G)** Expression of selected genes in time in mouse (solid orange line) and human differentiations (blue solid line) alognside with the rescaled expression in mouse (dashed orange line) using the estimated time factors for specific genes.

**Figure S2.**
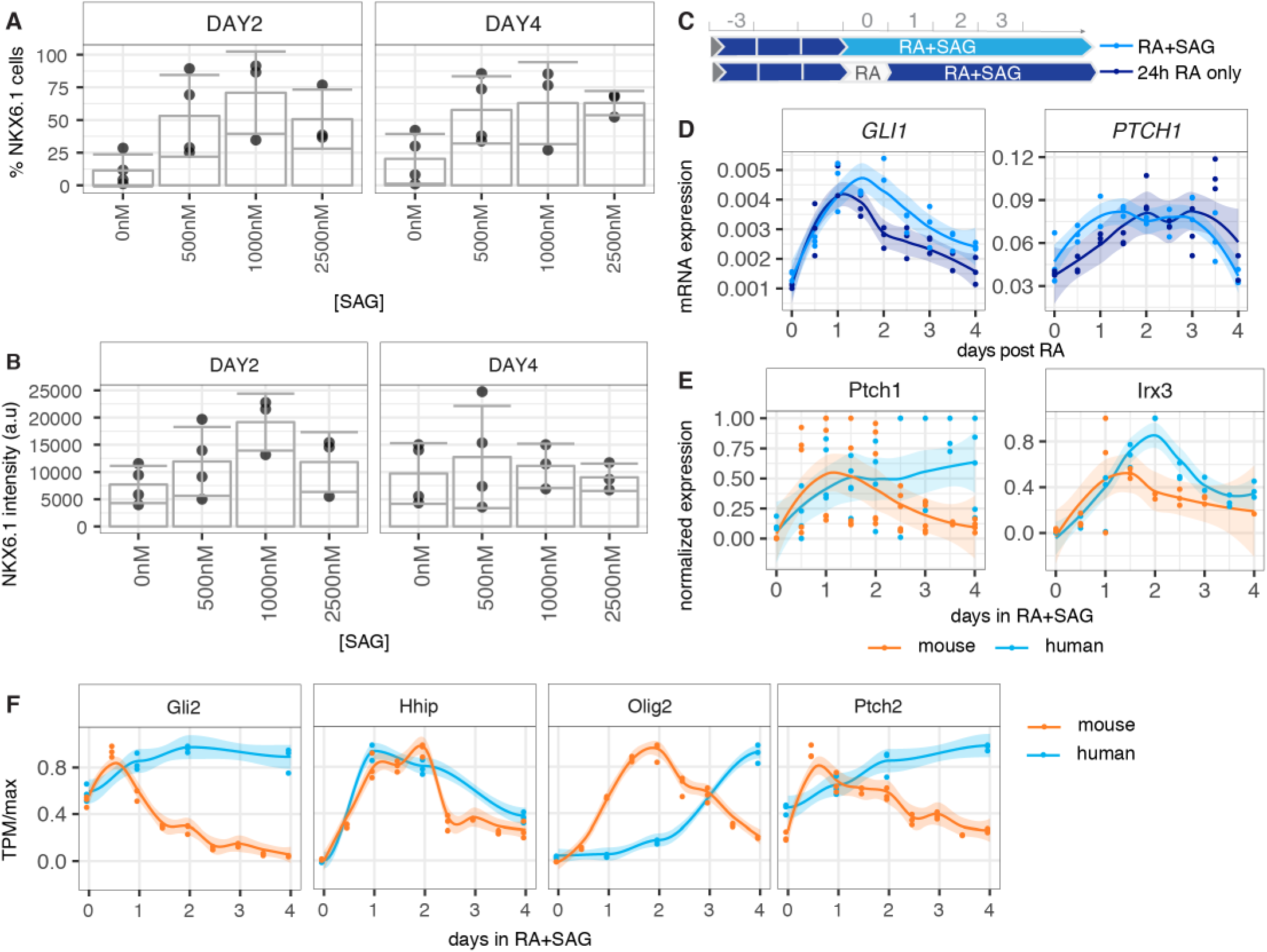
Dynamics of Shh at the time of induction in mouse and human neural progenitors. **(A)** Percentage of NKX6.1 measured by flow cytometry in SOX2-e×pressing cells at day 2 and day 4 after addition of SAG at increasing concentrations. **(B)** NKX6.1 mean intensity level from the same samples with increasing concentrations of SAG are comparable at day 2 and day 4. **(C)** Scheme outlining the standard differentiation protocol where RA and SAG are added at the same time (light blue), versus a treatment where SAG addition is delayed for 24h (dark blue). **(D)** RT-qPCR data reveals no substantial differences in the induction dynamics of Gli1, Ptchi from the moment of addition of SAG. **(E)** RT-qPCR data measured in 12h intervals reveals similar gene expression dynamics in mouse (orange) and human (blue) Ptchi and distinct Irx3 expression. **(F)** Normalized expression of other Hedghog target genes from the RNAseq. (Transcripts per million (TPM)).

**Figure S3.**
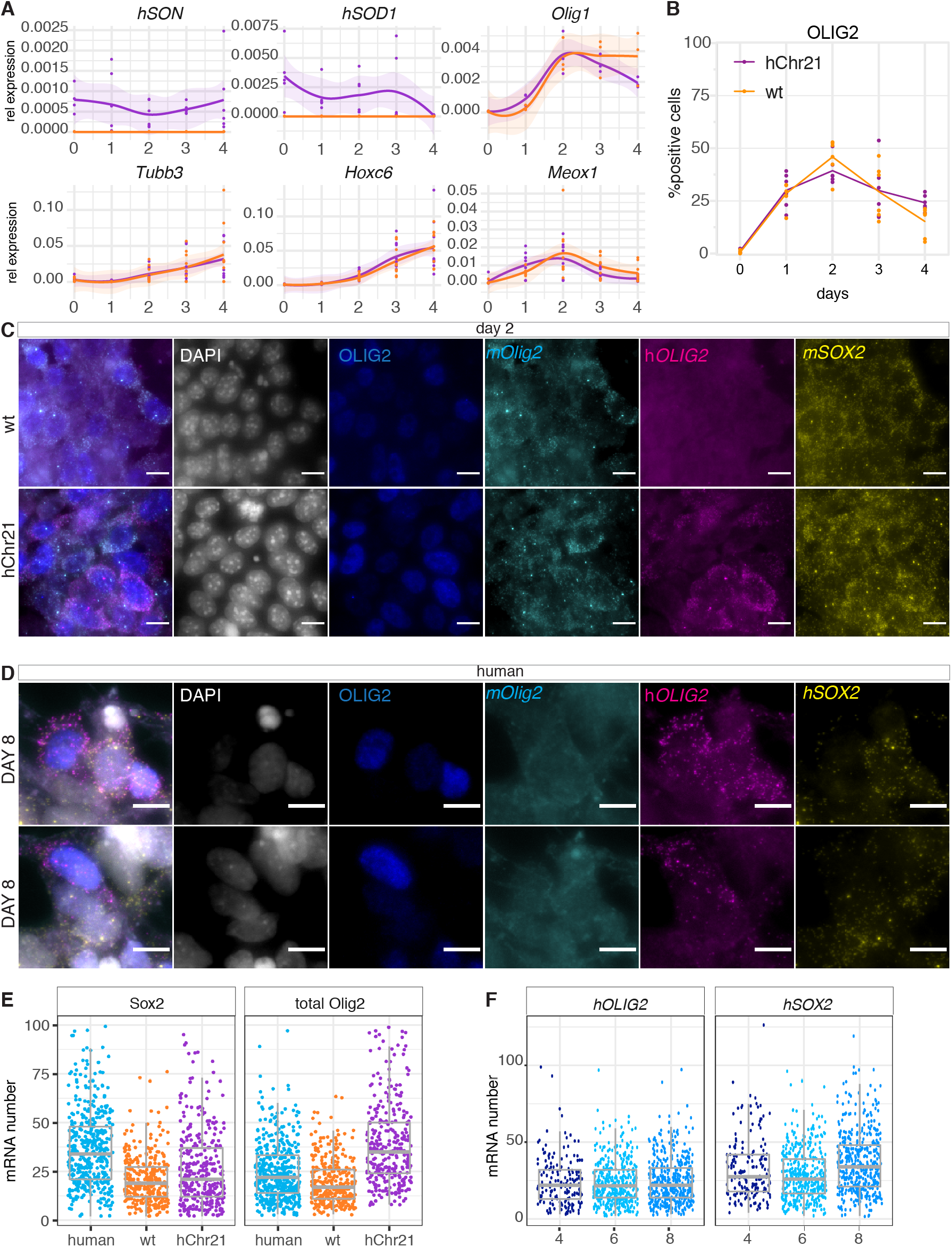
Characterization of mouse in vitro differentiations in the mouse ES cell line carrying hChr21. **(A)** Expression of Sod1 and Son human genes located on Chromosome 21, the neural markers Tubb3 and Olig 1 and the mesoderm marker Meoxl in wt and hChr21 differentiations. **(B)** Percentage of OLIG2 positive cells from days 0 to 4 in wt and hChr21 differentiation show no major changes in cell proportions over time. **(C)** Representative images of smFISH used for segmentation and quantification of wt and hChr21 mouse cells. **(D)** Representative images of smFISH used for segmentation and quantification of human MN differentiation. **(E)** Sox2 and total Olig2 mRNA number in human, wt and hChr21 neural progenitors. **(F)** *hSOX2* and *hOLIG2* mRNA number in human neural progenitors at days 4, 5 and 6. Scale bars 50μM.

**Figure S4.**
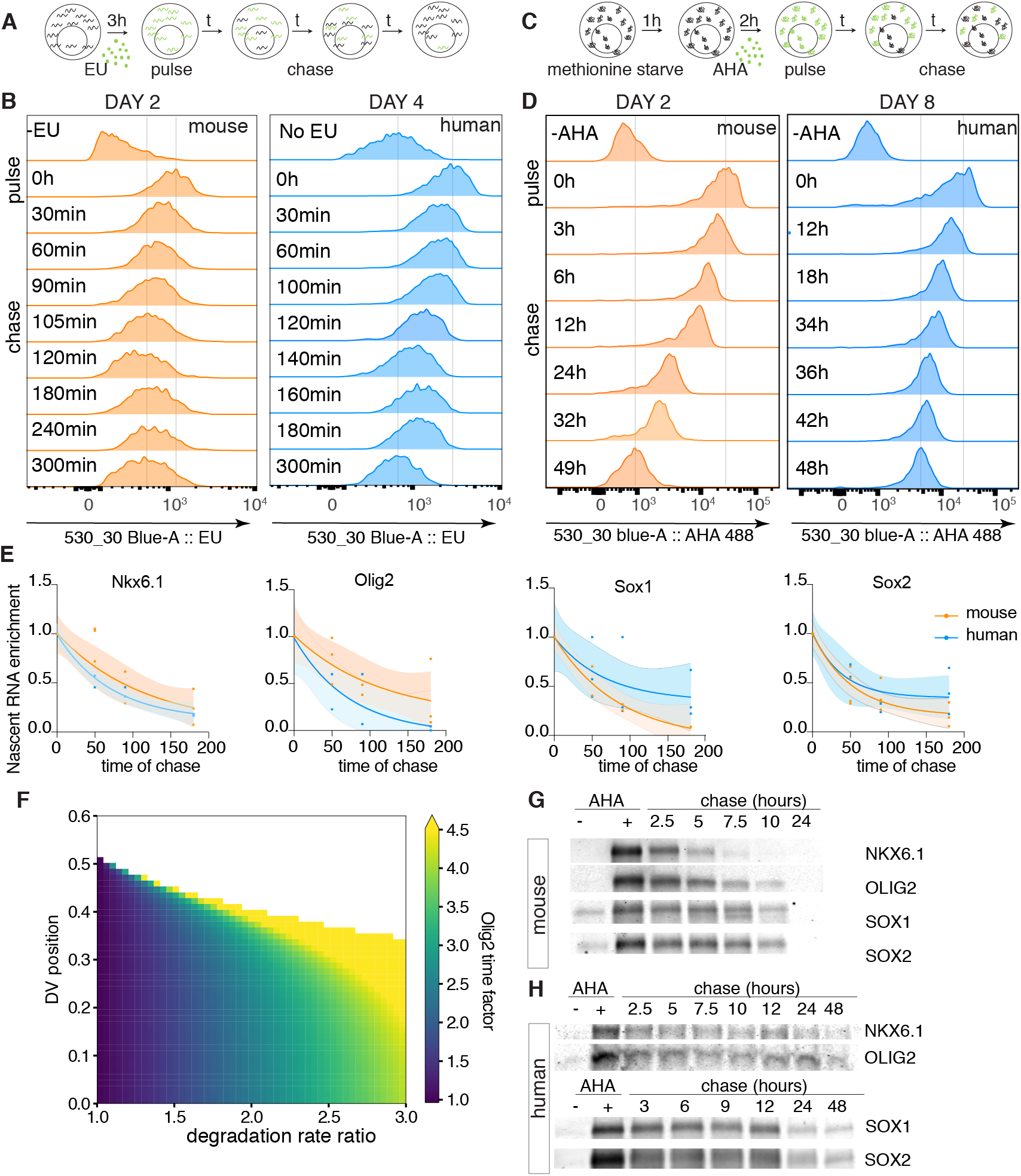
Protein stability in the GRN accounts for the speed differences between species. **(A)** Schematic of EU experiment to measure RNA half-lives in mouse and human neural progenitors. Cells are cultured for 3h with the uriridine analog EU (green). A sample that contains maximal levels of EU incorpotation is collected (pulse). Subsequently, EU is removed and cells are collected at various time points. **(B)** Representative histograms of EU incorporation intensity measurements to estimate mRNA half-life from mouse day 2 (orange) and human day 4 (blue) neural progenitors. **(C)** Schematic of AHA experiment to measure protein half-lives in mouse and human neural progenitors. Cells are grown on methionine-free media for 45h and then cultured for 2h with the methionine analog AHA (green). A sample that contains maximal levels of AHA incorpotation is collected (pulse). Subsequently, AHA is removed and cells are collected at various time points. **(D)** Representative histograms of AHA incorporation intensity measurements to estimate protein half-life from mouse day 2 (orange) and human day 8 (blue) neural progenitors. **(E)** Normalized RT-qPCR analysis of RNA enrichment relative to Actin of EU-nascent RNA captured on magenetic beads after pulse and chase experiments. **(F)** Estimated time factor as a function of the degradation rate ratio accros various dorsoventral (DV) positions. **(G)** Representative mouse **(G)** and human **(H)** detection of NKX6.1, OLIG2, S0X1 and SOX2 after AHA pulse-chase experiments on nascent proteins.

**Figure S5.**
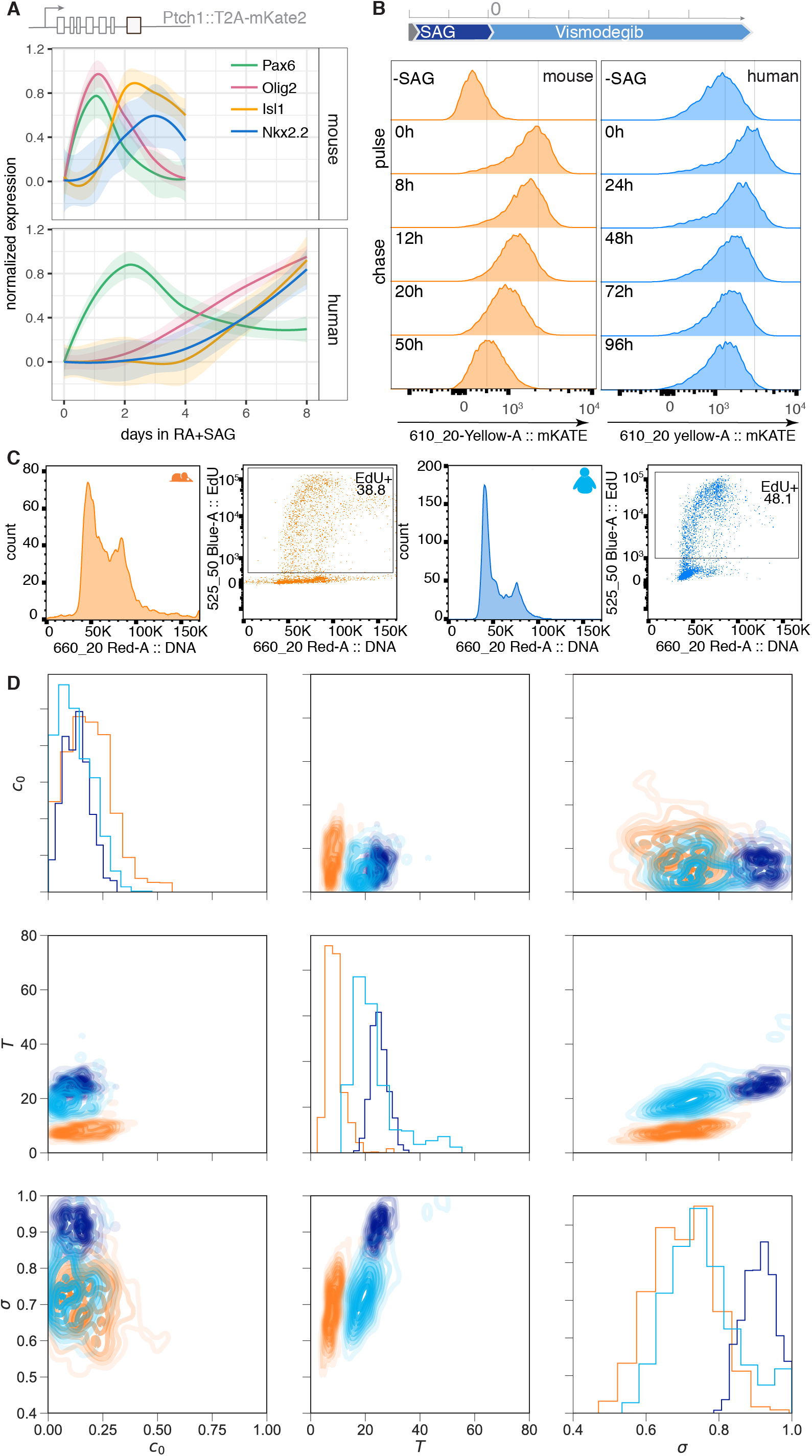
Protein stability and cell cycle measurements in mouse and human neural progenitors. **(A)** RT-qPCR analysis of Pax6, Olig2, Nkx2.2 and Isl1 expression in mouse and human differentiations from the Ptchl-mKATE2 targeted stem cell lines. **(B)** Representative histograms of mKATE2 intensity measurements to estimate protein half-life from mouse (orange) and human (blue) neural progenitors. **(C)** Representatve histograms of DNA content of mouse and human cells, and dual parameter plots of Alexa Fluor 488-conjugated EdU and DNA indicating the gates to quantify EdU positive cells. **(D)** Pairwise relationships between proportion of S phase estimations (c_0_), cell cycle lenght (*T*) and saturation (σ) in mouse and human cells.

